# Virological characteristics correlating with SARS-CoV-2 spike protein fusogenicity

**DOI:** 10.1101/2023.10.03.560628

**Authors:** MST Monira Begum, Kimiko Ichihara, Otowa Takahashi, Hesham Nasser, Michael Jonathan, Kenzo Tokunaga, Isao Yoshida, Mami Nagashima, Kenji Sadamasu, Kazuhisa Yoshimura, The Genotype to Phenotype Japan (G2P-Japan) Consortium, Kei Sato, Terumasa Ikeda

## Abstract

The severe acute respiratory syndrome coronavirus (SARS-CoV-2) spike (S) protein is essential in mediating membrane fusion of the virus with the target cells. Several reports demonstrated that SARS-CoV-2 S protein fusogenicity is reportedly closely associated with the intrinsic pathogenicity of the virus determined using hamster models. However, the association between S protein fusogenicity and other virological parameters remains elusive. In this study, we investigated the virological parameters of eleven previous variants of concern (VOCs) and variants of interest (VOIs) correlating with S protein fusogenicity. S protein fusogenicity was found to be strongly correlated with S1/S2 cleavage efficiency and plaque size formed by clinical isolates. However, S protein fusogenicity was less associated with pseudoviral infectivity, pseudovirus entry efficiency, and viral replication kinetics. Taken together, our results suggest that S1/S2 cleavage efficiency and plaque size could be potential indicators to predict the intrinsic pathogenicity of newly emerged SARS-CoV-2 variants.

## Introduction

Severe acute respiratory syndrome coronavirus 2 (SARS-CoV-2) is the causative agent of coronavirus disease 2019 (COVID-19). Since the first cases of a novel coronavirus infection were detected in Wuhan, Hubei Province, China, in December 2019^1,2^, the disease spread rapidly over the world and World Health Organization (WHO) declared it a Public Health Emergency of International Concern (PHEIC) as a COVID-19 pandemic on January 30, 2020^3^. Although WHO announced the end of PHEIC on May 5, 2023^4^, the COVID-19 pandemic is not over, thus far causing millions of deaths globally^5^.

SARS-CoV-2 displays a long RNA genome of approximately 30 kbp^6,7^, encoding 4, structural proteins [Spike (S), Envelope (E), Nucleocapsid (N), Membrane (M)], 9 accessory proteins (ORF3a, 3b, 6, 7a, 7b, 8, 9b, 9c, and 10), and 16 nonstructural proteins (NSPs; NSP1–16, encoded by the *ORF1a* and *ORF1b* genes), respectively^7–9^. SARS-CoV-2 S protein fulfills an important role in mediating the virus-target cell membrane fusion, thereby triggering viral entry into the target cells^10,11^. After its translation, cellular proteases (e.g., Furin in the Golgi apparatus of the infected cells) cleave the S protein into two subunits, S1 and S2^10^. This S1/S2 cleavage is important for SARS-CoV2 pathogenicity as SARS-CoV-2 lacking the furin cleavage site in the S protein exhibits attenuated viral pathogenicity in cell line, mouse, and hamster models^12^. The S protein is assembled as a trimer, inserted into viral particles along with the other viral components. The S1 and S2 subunits bind noncovalently and are exposed on the viral surface until meeting the target cells. To enter the target cells, the viral S protein and the host cell receptor angiotensin-converting enzyme 2 (ACE2) have to interact^13^. The receptor-binding domain engagement in the S1 subunit with ACE2 induces conformational changes in the S protein, leading to the S2’ site cleavage in the S2 subunit and fusion peptide insertion into the target cell membrane^10^. Next, the 6-bundle helix is formed by heptad repeats (HR) 1 and 2 of the S2 subunit as an indispensable fusion step, creating a fusion pore that facilitates genetic material transfer into the host cells. The S2’ site cleavage in the S2 subunit depends on the following SARS-CoV-2 entry routes^10^. First, the endosome-mediated entry pathway. Upon the S1 subunit-ACE2 interaction, a virus-ACE2 complex is internalized into the target cells by forming an endosome, where cathepsin L cleaves the S2 subunit S2’ site^10^. Second, the target cell surface, using transmembrane protease serine 2 (TMPRSS2) for the S2 subunit S2’ site cleavage^10,14^.

SARS-CoV-2 is continually evolving, with mutations appearing in its RNA genome since its discovery in 2019^6,15^. These *S* gene mutations influence transmissibility, pathogenicity, and vaccine- and viral infection-induced immune response resistance^11,14,16–19^. Multiple SARS-CoV-2 variants have emerged during the pandemic, certain among them predominantly spreading across the world. Initially, the Wuhan-Hu1 strain was outcompeted by the B.1 lineage (harboring the D614G mutation in the S protein), a potential ancestor of all recent variants^11,15,17^. Then, WHO defined the Alpha, Beta, Gamma, Delta, and Omicron BA.1, BA.2, and BA.5 variants as variants of concern (VOCs) as well as Lambda and Mu as variants of interest (VOIs)^11,15,17^, being already de-escalated from the VOC and VOI lists.

Efforts to describe the SARS-CoV-2 S protein features revealed that the S protein fusogenic potential of the SARS-CoV-2 variants is closely associated with their pathogenicity, determined using a hamster model without immunity against vaccines and viral infection (hereafter referred to as intrinsic pathogenicity)^20–24^. For example, the Delta variant possesses relatively more fusogenic S protein than the Omicron BA.1 and B.1.1 variants^21,23^. SARS-CoV-2 variants display higher intrinsic pathogenicity associated with the S protein-mediated fusogenicity strength^20–24^. However, the association of S protein-mediated fusogenicity with other virological parameters remains to be fully determined. In this study, we investigate the virological parameters of eleven SARS-CoV-2 variants correlating with S protein fusogenicity.

## Results

### S protein fusogenicity of eleven SARS-CoV-2 variants

SARS-CoV-2 S protein mediates the virus-infected cell membrane fusion^10,11^. To quantitatively monitor the fusion kinetics between effector cells expressing S protein from eleven SARS-CoV-2 variants (including the previous VOCs and VOIs) and target cell membranes, we performed S protein-mediated membrane fusion assay in Calu-3/DSP_1-7_ cells^20,22–27^. Although the Alpha, Beta, Gamma, Delta Lambda, Mu, and BA.5 S protein expression levels were lower than that of the B.1.1 S protein (harboring the D614G mutation) on the transfected HEK293 cell surface, the remaining S protein levels were comparable (**Fig. 1A**). Compared to B.1.1 S protein, the Wuhan S protein exhibited lower fusogenicity, while that related to the Alpha, Beta, Gamma, Delta, Lambda, and Mu S proteins was significantly higher (**Fig. 1B**). Notably, the Delta S protein exhibited profound fusogenicity^21,23^, being the highest of all tested S proteins (**Fig. 1B**). The BA.1 and BA.2 S proteins exerted lower fusogenicity compared to the B.1.1 S protein while that of the BA.5 S protein was not significantly different (**Fig. 1B**). These data indicate that SARS-CoV-2 S protein fusogenicity is different before and after Omicron variant emergence.

**Fig. 1.**
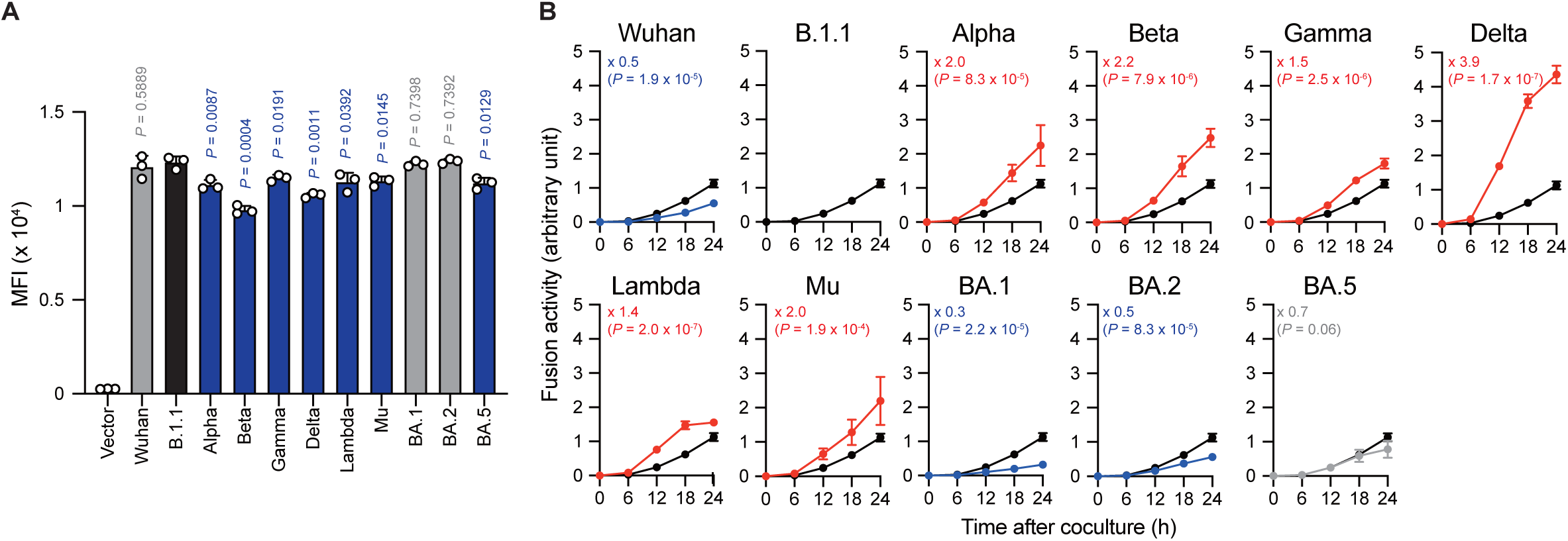
S protein fusogenicity correlates with S1/S2 cleavage efficiency. (**A)** Surface S protein expression in transfected HEK293 cells. The data represent the mean fluorescence intensity (MFI) with the mean ± standard deviation (SD) from 3 independent experiments. The statistical significance was tested against B.1.1 (black bar) using two-sided unpaired *t*-test. The P-values are indicated above each bar. The gray and blue bars indicate no significance and significantly reduced expression levels, respectively. (**B**) Indicated S variant fusion activities. The S protein-mediated membrane fusion assay was performed in Calu-3/DSP_1-7_ cells. The data at each time point are represented as the mean ± SD from 4 independent experiments. The numbers in each graph indicate the fold change 24 hours after S protein-expressing HEK293 and Calu-3 cell coculture compared to those expressing the B.1.1 S protein (black line). Statistical differences between B.1.1 S and each S variant across timepoints determined by multiple regression. The P-values are indicated in blue (reduction), red (increase), or gray (not significant) in each graph. We excluded 0-h data from the analyses. We indicated the Family-wise error rates (FWERs), calculated using the Holm method, in the figures.

### Correlation of S1/S2 cleavage efficiency with S protein fusogenicity

The SARS-CoV-2 S protein S1/S2 cleavage is pivotal, affecting viral fusogenicity^28^. For example, the level of Delta S protein S1/S2 cleavage is higher than that of the B.1.1 and BA.1 S proteins, being associated with higher fusogenic potential in the Delta variant^21,23,25,29^. To investigate whether the relationship between S protein S1/S2 cleavage and its fusogenicity could apply to S proteins from the other SARS-CoV-2 variants, we subjected HEK293 cells expressing each S protein to western blotting and quantified the full-length S and cleaved S2 band intensities (**Fig. 2A, 2B**). Full-length S and S2 band intensity variations indicated that each S protein exhibits a different susceptibility to cellular protease-induced S1/S2 cleavage. Consistent with previous results^21,23,25^, the Delta and Lambda S proteins displayed the highest S1/S2 cleavage efficiencies (**Fig. 2A, 2B**). In addition, the Wuhan, BA.1, BA.2, and BA.5 S protein S1/S2 cleavage was significantly lower than that of the B.1.1 S protein (**Fig. 2A, 2B**). Finally, we investigated how fusogenicity and S1/S2 cleavage efficiency correlate among the eleven SARS-CoV-2 variants (**Fig. 2C**). Remarkably, the correlation coefficient (R^2^ = 0.5975) indicated a strong positive correlation between fusogenicity and S1/S2 cleavage efficiency of all eleven SARS-CoV-2 S protein variants (**Fig. 2C**). In summary, these results demonstrated that SARS-CoV-2 S protein fusogenicity is strongly associated with S protein S1 and S2 subunit cleavage.

**Fig. 2.**
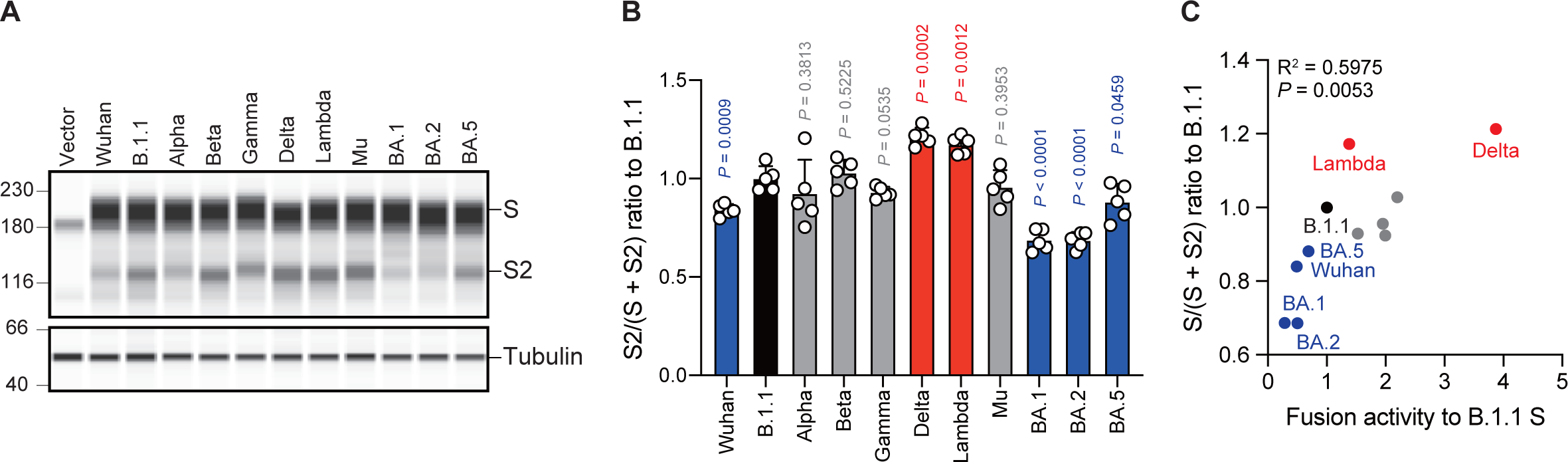
S protein fusogenicity correlates with S1/S2 cleavage efficiency. (**A**) Western blot analysis of S protein expressed in transfected HEK293 cells. The S and S2 proteins were detected using a rabbit anti-SARS-CoV-2 S2 polyclonal antibody. We used anti-α-tubulin as a loading control. (**B**) Indicated S variant S/S2 cleavage efficiencies. Each bar represents the S/(S + S2) values with the mean ± standard deviation from 5 independent experiments. The statistical significance was tested against B.1.1 (black bar) using two-sided unpaired *t*-tests. The P-values above each bar indicated in blue (reduction), red (increase), or gray (not significant). (**C**) Correlation between the S variant S/S2 cleavage efficiencies and fusion activities 24 hours after coculture. R^2^ denotes the correlation coefficient.

### Correlation of infection-induced plaque size with S protein fusogenicity

The Delta variant with higher fusogenic S protein levels reportedly forms larger plaques than the B.1.1 variant in infected VeroE6/TMPRSS2 cells^21^. However, the BA.1 variant encoding the lower fusogenic *S* gene displays smaller plaques compared to the B.1.1 variant^23^. These results suggest that S protein fusogenicity is potentially associated with viral infection, indicated the size of the plaques formed by clinical isolates. To address this question, we performed plaque assays using the eleven clinical isolates including the previous VOCs and VOIs and compared the plaque diameters formed by each clinical isolate in VeroE6/TMPRSS2 cells (**Fig. 3A, 3B**). We observed that the Wuhan, Alpha, Beta, Gamma, Delta, Lambda, and Mu variants formed significantly larger plaques than B.1.1 (**Fig. 3A, 3B**). In contrast, the BA.1, BA.2, and BA.5 variant-related plaque sizes were smaller compared to that of the B.1.1 variant (**Fig. 3A, 3B**). Interestingly, the SARS-CoV-2 infection-induced plaque size in VeroE6/TMPRSS2 strongly and positively correlated with S protein fusogenicity (**Fig. 3C**; R^2^ = 0.6982). These results indicate that SARS-CoV-2 S protein fusogenicity strongly influences the SARS-CoV-2 infection plaque size.

**Fig. 3.**
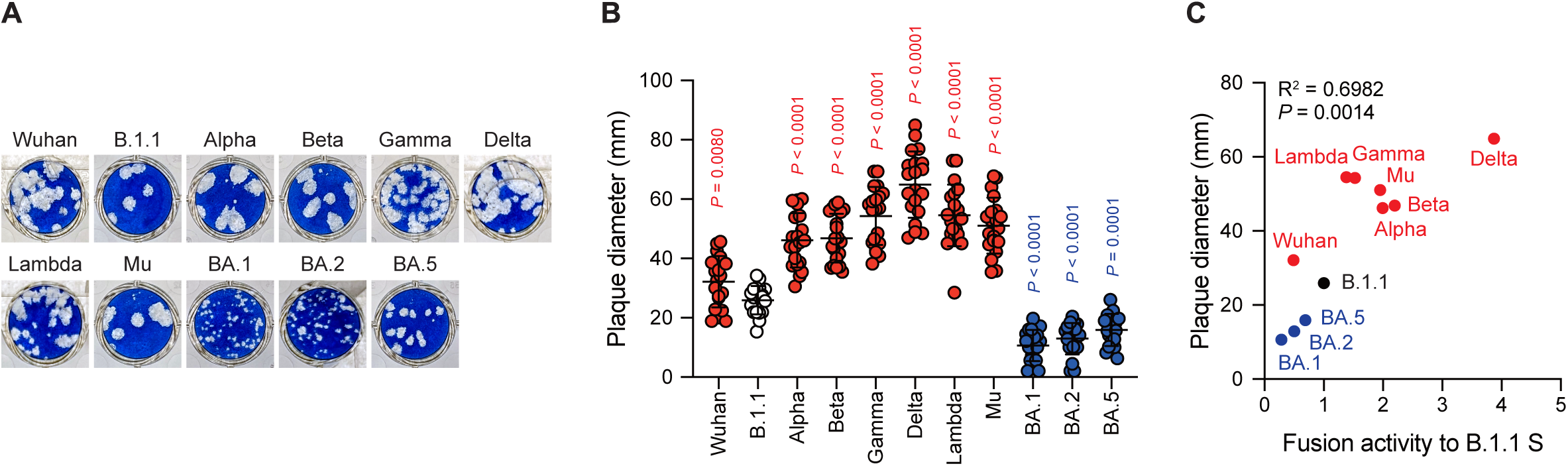
Clinical isolate-formed plaque size correlates with S protein fusogenicity. (**A**) Representative images of the plaque assays. (**B**) Plaque diameter summary. The data are represented as the plaque diameter with the mean ± standard deviation from 20 plaques per each variant. The statistical significance was tested against B.1.1 (white circles) by using two-sided unpaired *t*-tests. The P-values above each bar are indicated in red (increase) or blue (reduction). (**C**) Correlation between the plaque diameter and fusion activity of the corresponding variants 24 hours after coculture. R^2^ denotes the correlation coefficient.

### Correlation of SARS-CoV-2 S pseudovirus infectivity with S protein fusogenicity

Next, we examined the correlation of S protein-mediated cell-free viral infection with S protein fusogenicity. We used HEK293T cells to produce HIV-1 virions pseudotyped with SARS-CoV-2 S protein carrying the C-terminal deletion of 19 amino acids (Δ19CT), normalized by the p24 concentration, and measured viral infectivity as described previously^20–24,26,27,29–31^. Our results revealed that the Delta and BA.5 S pseudoviral infectivity measured using the HOS-ACE2/TMPRSS2 system were significantly higher than that of the B.1.1 S pseudovirus while that of the Wuhan, Alpha, Beta, Lambda, Mu, BA.1 and BA.2 S pseudoviruses significantly decreased in HOS-ACE2/TMPRSS2 cells compared to B.1.1 (**Fig. 4A**). We observed no statistically significant difference between the Gamma S pseudoviral infectivity and that of the B.1.1 S pseudovirus (**Fig. 4A**). Notably, we noticed that the infectivity degrees of the eleven SARS-CoV-2 variant-derived S protein-mediated cell-free pseudoviruses did not correlate with the fusogenicity mediated by these S proteins (**Fig. 4B**, R^2^ = 0.0033). This discrepancy suggests that the each S protein incorporation level into the pseudoviruses might vary, thereby leading to incorrect viral infectivity validation. To test this hypothesis, we analyzed the S protein incorporation levels in the viral particles using western blotting (**Fig. 4C, 4D**). The S protein band intensity quantification revealed that compared to the B.1.1 S pseudovirus, Wuhan, Alpha, Mu, BA.1, BA.2, and BA.5 variant-derived S proteins incorporated less in the pseudovirus particles (**Fig. 4C, 4D**). These data indicate variable S protein incorporation levels, prompting that viral infectivity per incorporated S protein should be evaluated. Although most SARS-CoV-2 S pseudoviral infectivity divided by S protein incorporation levels became comparable or lower than that of the B.1.1 S pseudovirus, only those of the BA.2 and BA.5 S pseudoviruses were significantly higher (**Fig. 4E**). In addition, the R^2^ value between the cell-free pseudoviral infectivity per S protein and fusogenicity mediated by these S proteins slightly changed from 0.0033 to 0.1482 (**Fig. 4B, 4F**). In summary, these data suggest that S protein fusogenicity might have little effect on pseudovirus infection in target cells.

**Fig. 4.**
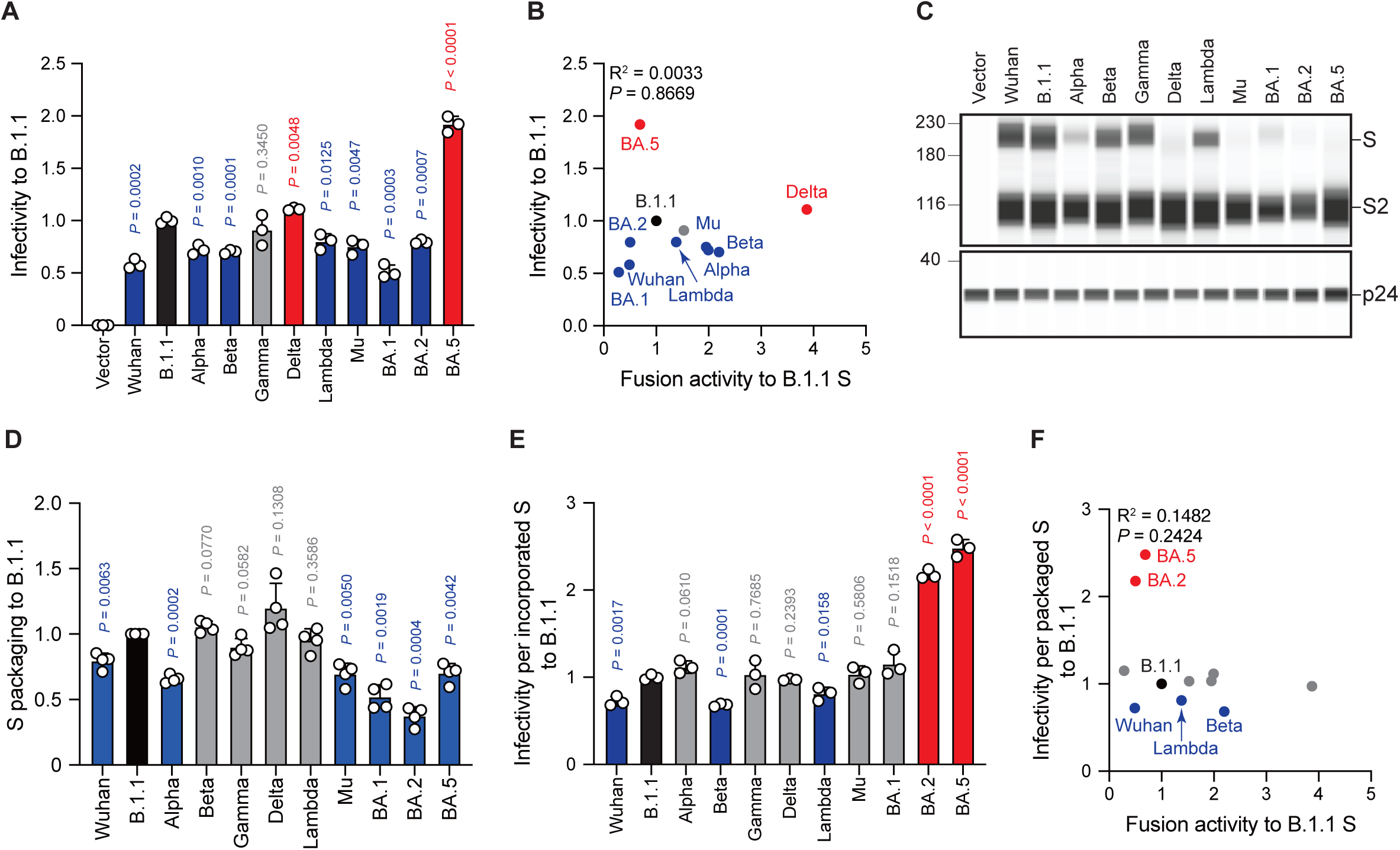
SARS-CoV-2 S pseudovirus infectivity does not correlate with S protein fusogenicity. (**A**) Representative SARS-CoV-2 pseudoviral infectivity data. We used HEK293T cells to produce the HIV-1-based pseudoviruses with the indicated SARS-CoV-2 S proteins for HOS-ACE2/TMPRSS2 cell infection. We measured the intracellular luciferase activity of each virus-infected cell and expressed as viral infectivity relative to that of B.1.1 S pseudovirus (black bar) with the mean ± standard deviation (SD) from three independent experiments. The statistical significance was tested against B.1.1 S using two-sided unpaired *t*-tests. The P-values above each bar are indicated in blue (reduction), red (increase), or gray (not significant). (**B**) Correlation between relative pseudoviral infectivity and fusion activity 24 hours after coculture. (**C**) Representative western blot data of S proteins incorporated in the pseudoviruses. S and S2 were detected using a rabbit anti-SARS-CoV-2 S2 polyclonal antibody. We used p24 as a loading control. (**D**) S protein incorporation levels in the pseudoviruses relative to B.1.1. The data are represented as the mean ± SD from 4 independent experiments. The statistical significance was tested against B.1.1 S (black bar) using two-sided paired *t*-tests. The P-values above each bar are indicated in blue (reduction) or gray (not significant). (**E**) Representative infectivity data of SARS-CoV-2 pseudoviruses per S protein incorporation level. SARS-CoV-2 pseudoviral infectivities were divided by the incorporation level of each S protein and the viral infectivities were normalized by that of the B.1.1 S pseudovirus (black bar). The data are represented as the mean ± SD from three independent experiments. The statistical significance was tested against B.1.1 S using two-sided paired *t*-tests. The P-values above each bar are indicated in blue (reduction), red (increase), or gray (not significant). (**F**) Correlation between relative pseudoviral infectivity per S protein incorporation level and fusion activity 24 hours after coculture. R^2^ denotes the correlation coefficient in each panel where indicated.

### Correlation of SARS-CoV-2 S pseudovirus entry efficiency with S protein fusogenicity

The aforementioned pseudovirus assays indicated a weak correlation between S protein fusogenicity and SARS-CoV-2 S pseudoviral infectivity even after normalizing the S protein incorporation levels into the viral particles (**Fig. 4F**). These data suggest that HIV-1 infection mechanisms, such as reverse transcription and integration, might affect the pseudovirus assay results. To address this hypothesis, we performed a BlaM-Vpr assay using S protein with Δ19CT enabling us to quantify cell-free virus entry efficiency into HOS-ACE2/TMPRSS2 cells (**Fig. 5A**). The results indicated that all S pseudoviruses exhibited comparable or lower entry efficiencies compared to the B.1.1 S pseudovirus (**Fig. 5A**). Similar to the pseudovirus assays (**Fig. 4B, 4C**), we normalized the fluorescence signals indicating entry efficiency into the target cells by the S protein incorporation levels due to the variable S protein incorporation levels into the viral particles (**Fig. 5B-D**). Interestingly, although the entry efficiency determined by BlaM-Vpr assays and normalized by the viral S protein level strongly correlated with the S pseudovirus infectivities per S protein incorporation levels (**Fig. 5E**; R^2^ = 0.6209), the relationship with S protein fusogenicity was weak, consistently with pseudoviral infectivity (**Figs. 4F, 5F**; R^2^ = 0.1482 and 0.0014, respectively). These data support the results suggesting that the S protein fusogenic potential could affect little pseudovirus entry into the HOS-ACE2/TMPRSS2 cells.

**Fig. 5.**
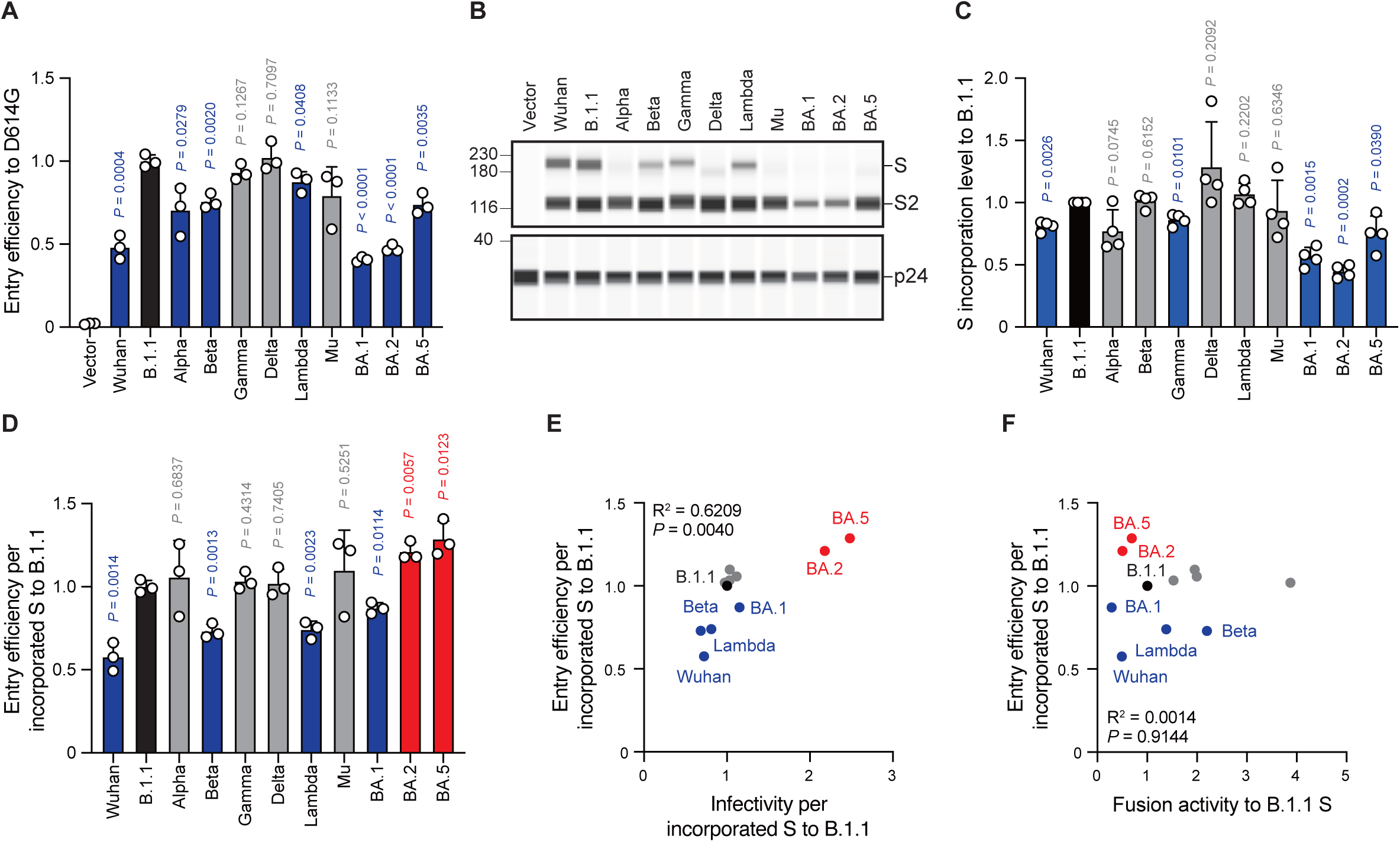
S protein-mediated pseudovirus entry efficiency does not correlate with S protein fusogenicity. (**A**) Representative BlaM-Vpr assay data. We used HEK293T cells to produce HIV-1-based pseudoviruses with the indicated SARS-CoV-2 S proteins for HOS-ACE2/TMPRSS2 cell infection. The cleaved CCF4 signal ratio to the sum of uncleaved and cleaved CCF4 signals was calculated and indicated as the entry efficiency relative to that of the B.1.1 S pseudovirus (black bar) with the mean ± standard deviation (SD) from three independent experiments. The statistical significance was tested against B.1.1 S using two-sided unpaired *t*-tests. The P-values above each bar are indicated in blue (reduction) or gray (not significant). (**B**) Representative western blot data of S proteins incorporated in the pseudoviruses. S and S2 were detected using a rabbit anti-SARS-CoV-2 S2 polyclonal antibody. We used p24 as a loading control. (**C**) S protein incorporation level in the pseudoviruses relative to B.1.1. The data are represented as the mean ± SD from 4 independent experiments. The statistical significance was tested against B.1.1 S (black bar) using two-sided paired *t*-tests. The P-values above each bar are indicated in blue (reduction) or gray (not significant). (**D**) Representative BlaM-Vpr data normalized by the S protein incorporation levels. S protein-mediated SARS-CoV-2 pseudovirus entry efficiencies were divided by each S protein incorporation level and the entry efficiencies were normalized by that of the B.1.1 S pseudovirus (black bar). The data are represented as the mean ± SD from three independent experiments. The statistical significance was tested against B.1.1 S using two-sided paired *t*-tests. the P-values above each bar are indicated in blue (reduction), red (increase), or gray (not significant). (**E**) Correlation between relative pseudoviral infectivities and S protein-mediated entry efficiencies per S protein incorporation levels. (**F**) Correlation between S protein-mediated entry efficiency and fusogenicity 24 hours after coculture. R^2^ denotes the correlation coefficient in each panel where indicated.

### Correlation of replication kinetics of clinical isolates in VeroE6/TMPRSS2 and Calu-3 cells with S protein fusogenicity

To investigate the clinical isolate replication kinetics, we inoculated the eleven SARS-CoV-2 variants into VeroE6/TMPRSS2 and Calu-3 cells and measured the viral RNA levels in the cell culture supernatants using RT–qPCR (**Fig. 6A, 6B**). Compared to the B.1.1 variant, the Lambda, BA.1, and BA.5 variants displayed slower replication kinetics in VeroE6/TMPRSS2 cells while the other variants were comparable (**Fig. 6A**). In Calu-3 cells, the Alpha, Gamma, Delta, Mu, and BA.5 variant replication kinetics indicated significantly higher and that of the BA.2 variant lower values than that of the B.1.1 variant (**Fig. 6B**). The other variants, such as Wuhan, Beta, Lambda, and BA.1, displayed no statistically significant difference (**Fig. 6B**). Finally, we aimed at addressing a potential correlation between viral RNA production in VeroE6/TMPRSS2 or Calu-3 cells 48 hours postinfection and S protein fusogenicity (**Fig. 6C, 6D**). However, we obtained low R^2^ values, indicating no and weak viral RNA production correlation with S protein fusogenicity (**Fig. 6C, 6D**; R^2^ = 0.0152 in VeroE6/TMPRSS2 cells and R^2^ = 0.1543 in Calu-3 cells, respectively). These data suggest that S protein fusogenicity might not or could only slightly affect viral RNA production during viral replication in VeroE6/TMPRSS2 and Calu-3 cells.

**Fig. 6.**
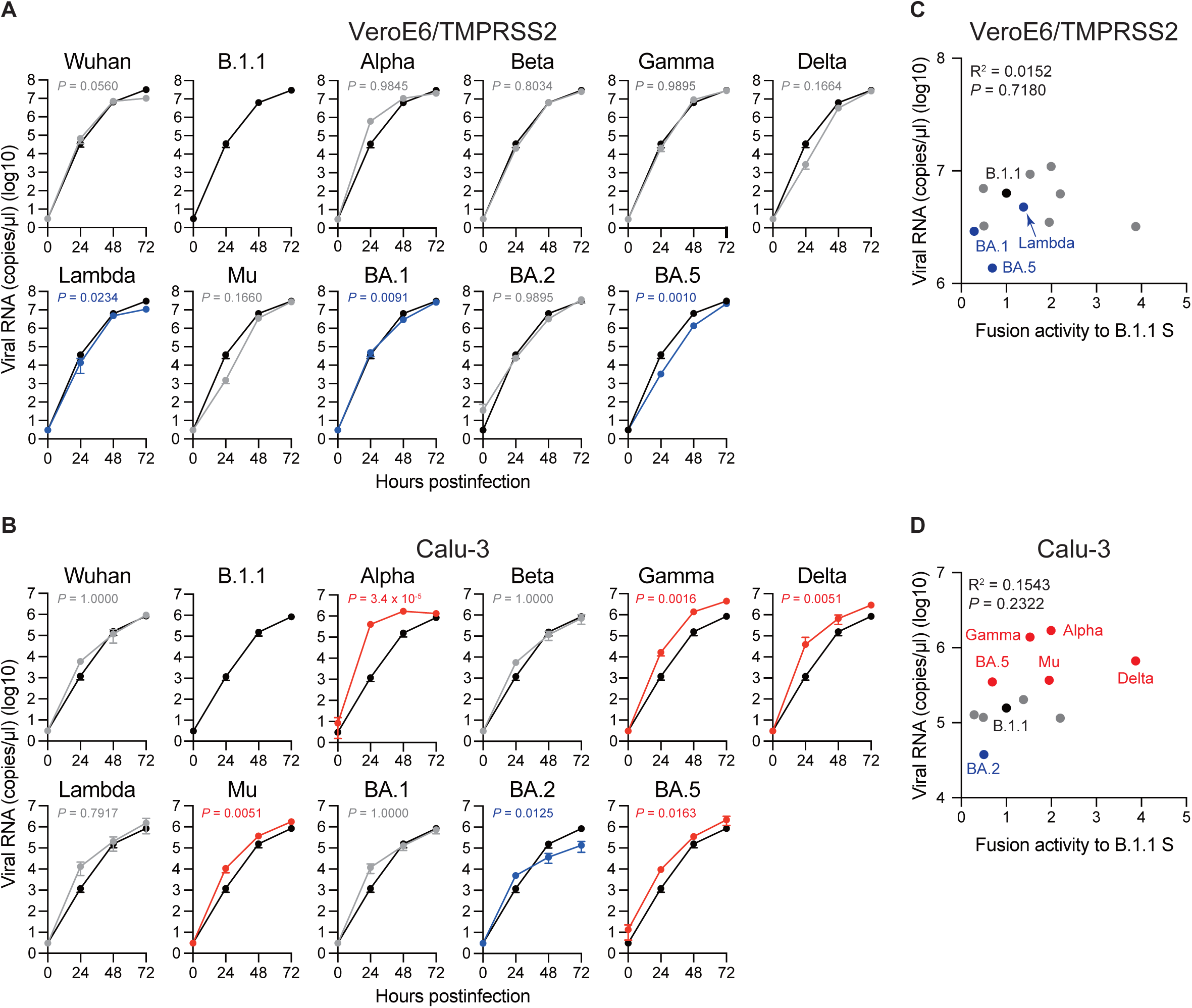
Replication kinetics of the eleven SARS-CoV-2 variants in VeroE6/TMPRSS2 and Calu-3 cells. (**A**) Representative replication kinetics of indicated clinical isolates in VeroE6/TMPRSS2 cells. The viral RNA copy number in the supernatant was quantified using RT–qPCR. The data at each time point are represented as the mean ± standard deviation (SD) from 4 independent experiments. The statistical differences between B.1.1 S (black line) and each S variant across the timepoints were determined using multiple regression. The P-values are indicated in blue (reduction) or gray (not significant) in each graph. The 0-h data were excluded from the analyses. (**B**) Representative replication kinetics of indicated clinical isolates in Calu-3 cells. The viral RNA copy number in the supernatant was quantified using RT–qPCR. The data at each time point are represented as the mean ± SD from 4 independent experiments. The statistical differences between B.1.1 S (black line) and each S variant across the timepoints were determined using multiple regression. The P-values are indicated in blue (reduction), red (increase), or gray (not significant) in each graph. The 0-h data were excluded from the analyses. We indicated the Family-wise error rates (FWERs), calculated using the Holm method. (**C**) Correlation between viral RNA copy numbers 48 hours postinfection and the fusogencity of the indicated S variants 24 hours after coculture. (**D**) Correlation between viral RNA copy numbers 48 hours postinfection and fusogenicities of the indicated S variants 24 hours after coculture. R^2^ denotes the correlation coefficient in each panel where indicated.

## Discussion

SARS-CoV-2 constantly evolves with mutations in its viral genome since its emergence in late 2019. In particular, the virological characteristics of the Omicron variants, such as transmissibility, pathogenicity, and immunity resistance, are rather different from those of the pre-Omicron variants^11,15,18,19,32,33^. The *S* gene is the most variable gene in the emerged SARS-CoV-2 variants. The SARS-CoV-2 S protein is pivotal in mediating the virus-target cell membrane fusion. Previous studies indicated that the SARS-CoV-2 S protein fusogenicity is closely associated with intrinsic pathogenicity^20–24^. Therefore, describing S protein features is of utmost importance. In this study, we investigated the correlation between S protein fusogenicity and other virological parameters in eleven SARS-CoV-2 variants including previous SARS-CoV-2 VOCs and VOIs. The S protein fusogenicity strongly correlated with S protein S1/S2 cleavage in transfected HEK293 cells (**Fig. 2C**) and the plaque size in VeroE6/TMPRSS2 cells infected by clinical isolates (**Fig. 3C**). However, the S protein fusogenicity was weakly associated with S protein-mediated pseudoviral infectivity (**Figs. 4B, F**) and entry efficiency in HOS-ACE2/TMPRSS2 cells (**Fig. 5F**) as well as viral replication kinetics in VeroE6/TMPRSS2 and Calu-3 cells (**Figs. 6C, D**). Taken together, our data suggest that, similar to SARS-CoV-2 S protein fusogenicity, S1/S2 cleavage and plaque size could be potential indicators to predict the intrinsic pathogenicity of newly emerged SARS-CoV-2 variants.

Newly emerged SARS-CoV-2 variant pathogenicity is rather different in clinical settings before and after the emergence of the Omicron BA.1 variant^18^. Although the Delta variant resulted in more severe outcome in infected patients, those of the Omicron BA.1 variant were attenuated^18^. Newly emerged SARS-CoV-2 variant pathogenicity might be predictable by measuring S protein fusogenicity as the latter is reportedly closely associated with the intrinsic pathogenicity of certain SARS-CoV-2 variants^20–24^. For example, the Delta variant displays more fusogenic S proteins and represents higher pathogenicity as demonstrated in a hamster model^21^. In contrast, the S protein of the Omicron BA.1 variant is less fusogenic and its pathogenicity also remains relatively low^23,24^. In support of these clinical and experimental observations, our data demonstrated that the S proteins of six pre-Omicron variants (Alpha, Beta, Gamma, Delta, Lambda, and Mu) exhibited higher fusogenicity than the D614G S protein based on our S protein-mediated membrane fusion assay, while the Omicron S proteins of the BA.1, BA.2 and BA.5 variants yielded comparable or lower fusogenicities (**Fig. 1B**). Moreover, we demonstrated that the S protein S1/S2 cleavage efficiency and plaque size in clinical isolates correlated with S protein fusogenicity (**Figs. 2C, 3C**), suggesting that these parameters could serve as potential markers for intrinsic pathogenicity prediction of newly emerged SARS-CoV-2 variants.

Nevertheless, the relationship between viral fusogenicity and viral intrinsic pathogenicity is not applied to recently emerged Omicron subvariants, the BQ.1.1 and XBB.1 variants^26,27^. Although these variants demonstrated higher fusogenicity than the BA.5 and BA.2.75 variants, their intrinsic pathogenicity was comparable or lower^26,27^. These unexpected results might be explained by a scenario where non-S proteins could cancel the S protein-induced viral intrinsic pathogenicity increase. In fact, *ORF1ab*^34–37^, *ORF3a*^38–40^, *ORF6*^38,41^, *ORF7a*^42^, and *ORF8*^43–45^ gene deletions or mutations and those of genes downstream of the *S* gene^46^ are reportedly associated with viral growth attenuation in cell lines or pathogenicity in infected animal models. Remarkably, the BQ.1.1 and XBB.1 variants display at least six and seven amino acid substitutions in the non-S protein coding region compared to the BA.5 and BA.2.75 variants^26,27^. One or some of these substitutions might be involved in attenuating the S protein-augmented intrinsic pathogenicity. Further studies would be required to investigate the non-S gene-intrinsic pathogenesis interaction.

SARS-CoV-2 S protein fusogenicity is essential for viral entry into the target cells. Nevertheless, SARS-CoV-2 S pseudoviral infectivity, entry efficiency, and viral replication indicated less correlation with S protein fusogenicity (**Figs. 4F, 5F, 6C, 6D**). Concerning our pseudovirus and BlaM-Vpr assays, we used lentiviruses pseudotyped with different S proteins of interest. In addition, we used HOS-ACE2/TMPRSS2 cells as non-natural SARS-CoV-2 target cells that stably express ACE2 and TMPRSS2. The related observations could contribute to drawing a difference from viral infection *in vivo*. Furthermore, viral RNA production during viral replication in VeroE6/TMPRSS2 and Calu-3 cells did not correlate with S protein fusogenicity. Indeed, various studies described that at least pseudoviral infectivity is not necessarily consistent with viral fusogenicity, pathogenicity, and epidemiology^22,24,26,27,44,47–50^. Accordingly, our pseudoviral infectivity, BlaM-Vpr, and viral replication assay-related data might be reasonable. However, performing these assays in at least primary lung cells would need to better understand how virological parameters (e.g., SARS-CoV-2 S pseudoviral infectivity, entry efficiency, and viral replication kinetics) could affect viral pathogenicity.

In summary, we revealed that S protein S1/S2 cleavage efficiency and clinical isolate plaque size are associated with SARS-CoV-2 S protein fusogenicity. Since the S protein fusogenicity is closely associated with intrinsic viral pathogenicity, these virological parameters could be used as potential markers to predict the intrinsic pathogenicity of newly emerged SARS-CoV-2 variants. However, the relationship between viral fusogenicity and intrinsic pathogenicity do not apply to the BQ.1.1 and XBB.1 variants. Therefore, further studies would be required to precisely describe the virological features of newly emerged SARS-CoV-2 variants, such as viral pathogenicity-determining factor(s).

## Methods

### Cell lines

HEK293T cells (a human embryonic kidney cell line; ATCC, CRL-1573), HEK293 cells (a human embryonic kidney cell line; ATCC, CRL-1573) and HOS-ACE2/TMPRSS2 cells (HOS cells stably expressing human ACE2 and TMPRSS2)^51^ were cultured in high glucose Dulbecco’s Modified Eagle Medium (DMEM, high glucose) (Wako, Cat# 044-29765) supplemented with 10% fetal bovine serum (FBS) (NICHIREI, Cat# 175012) and 1% penicillin/streptomycin (P/S) (Wako, Cat# 168-23191). VeroE6/TMPRSS2 cells (VeroE6 cells stably expressing human TMPRSS2; JCRB Cell Bank, JCRB1819)^52^ were maintained in DMEM (low glucose, Wako, Cat# 041-29775) containing 10% FBS, G418 (1 mg/ml; Wako, Cat#070-06803) and 1% P/S. Calu-3 cells (ATCC, HTB-55) were maintained in EMEM (Wako, Cat#055-08975) containing 20% FBS and 1% P/S. Calu-3/DSP_1-7_ cells (Calu-3 cells stably expressing DSP_1-7_)^53^ were maintained in EMEM (Wako, Cat# 055-08975) containing 20% FBS and 1% P/S. All cells were maintained at 37°C with 5% CO_2_.

### Virus preparation

SARS-CoV-2 Wuhan variant (strain SARS-CoV-2/Hu/DP/Kng/19-020 strain, Genbank accession no. LC528232)^54^ was provided by Drs. Tomohiko Takasaki and Jun-Ichi Sakuragi (Kanagawa Prefectural Institute of Public Health). SARS-CoV-2 Alpha (strain QHN001, GISAID ID: EPI_ISL_804007)^54^, Beta (strain TY8-612, GISAID ID: EPI_ISL_1123289)^55^, Gamma (strain TY7-503, GISAID ID: EPI_ISL_877769)^54,55^, Delta (strain TY11-927, GISAID ID: EPI_ISL_2158617), Lambda (strain TY33-456, GISAID ID: EPI_ISL_4204973), Mu (strain TY26-717, GISAID ID: EPI_ISL_4470503), Omicron BA.1 (strain TY38-873, GISAID ID: EPI_ISL_7418017)^23^, and Omicron BA.2 (strain TY40-385, GISAID ID: EPI_ISL_9595859)^22,27^ variants were obtained from National Institute of Infectious Diseases. D614G-bearing B.1.1 (strain TKYT41838, Genbank accession no. LC606020) and BA.5 (strain TKYS14631; GISAID ID: EPI_ISL_12812500)^20,22^ variants were provided by Tokyo Metropolitan Institute of Public Health.

Virus propagation was performed as previously described^27,54,56^. Briefly, VeroE6/TMPRSS2 cells (5 × 10^6^ cells) were seeded in a T-75 flask the day before infection. Virus was diluted in virus dilution buffer [1M HEPES, DMEM (low glucose), Non-essential Amino acid (gibco, Cat# 11140-050), 1% P/S] and the dilution buffer containing virus was added to the flask after removing the initial medium. After 1 hour of incubation at 37°C, the supernatant was replaced with 15 ml of 2% FBS/DMEM (low glucose) and cell culture was continued to incubate at 37°C until visible cytopathic effect (CPE) was clearly observed. Then, cell culture supernatant was collected, centrifuged at 300 × *g* for 10 minutes and frozen at –80°C as working virus stock. The titer of the prepared working virus was determined as the 50% tissue culture infectious dose (TCID_50_)^20,26,27^. The day before infection, VeroE6/TMPRSS2 cells (10,000 cells) were seeded in a 96-well plate and infected with serially diluted working virus stocks. The infected cells were incubated at 37°C for 4 days and the appearance of CPEs in the infected cells was observed by a microscope. The value of TCID_50_/ml was calculated by the Reed-Muench method^57^.

### Plasmid construction

Plasmids expressing the codon-optimized SARS-CoV-2 S proteins of Wuhan^30^, B.1.1 (*S* gene bearing D614G mutation)^30^, Alpha^21^, Beta^21^, Gamma^58^, Delta^21^, Lambda^58^, Mu^59^, BA.1^23^, BA.2^24^, and BA.5^20^ variants were prepared in previous studies. For creation of S proteins with 19 amino acids deletion at the C-terminal end, each *S* gene was amplified with primers; forward, 5’-NNN NNG GTA CCA TGT TTG TGT TCC TGG TGC T-3’ and reverse 5’-GTG GCG GCC GCT CTA GAT TCA ACA ACA GGA GCC ACA GGA A-3’. The resulting PCR fragment was digested with KpnI (New England Biolabs, Cat# R3142S) and NotI (New England Biolabs, Cat# R3189L) and inserted into the corresponding site of the pCAGGS vector^60^. All constructs were confirmed by Sanger sequencing (AZENTA) and the sequence data were analyzed by Sequencher v5.4.6 software (Gene Codes Corporation).

### SARS-CoV-2 S-based fusion assay

A SARS-CoV-2 S-based fusion assay with dual split protein (DSP) encoding *Renilla luciferase* (*RL*) and *GFP* genes was performed as previously described^25^. On day 1, effector cells (i.e., S-expressing) and target cells (Calu-3/DSP_1-7_ cells) were prepared at a density of 0.6-0.8 × 10^6^ cells/well in a 6 well plate. On day 2, to prepare effector cells, HEK293 cells were cotransfected with the S expression plasmids (400 ng) and pDSP ^61,62^ (400ng) using TransIT-LT1 (Takara, Cat# MIR2306). On day 3 (24 hours post transfection), effector cells were detached by pipetting and 16,000 effector cells/well were reseeded into a 96 well black plate (PerKinElmer, Cat# 6005225), and target cells were reseeded at a density of 1 × 10^6^ cells/2 ml/well in a 6 well plate. On day 4 (48 hours post transfection), the target cells were incubated with Enduren live cell substrate (Pomega, Cat# E6481) for 3 hours and then detached, and 32,000 target cells/well were added to a 96 well plate with effector cells. RL activity was measured at the indicated time points using a Centro XS3 LB960 (Berthhold Technologies). To measure the surface expression level of the S protein, effector cells were stained with rabbit anti-SARS-CoV-2 S S1/S2 polyclonal antibody (Thermo Fisher Scientific, Cat#PA5-112048, 1:100 dilution). Normal rabbit IgG (SouthernBiotech, Cat# 0111-01, 1:100) was used as negative control, and APC-conjugated goat anti-rabbit IgG polyclonal antibody (Jackson immunoresearch, Cat# 111-136-144, 1:50) was used as a secondary antibody. The expression level of the surface S proteins was detected by FACS Canto II (BD Biosciences) and analyzed by FlowJo software v10.7.1 (BD Biosciences) (**Fig. 1A**). RL activity was normalized to the mean fluorescence intensity (MFI) of surface S proteins, and the normalized values are shown as fusion activity (**Fig. 1B**).

### Pseudovirus assay

For pseudovirus production, lentivirus (HIV-1)-based luciferase-expressing reporter viruses were pseudotyped with the SARS-CoV-2 S protein^20,22,30,50^. HEK293T cells (2 × 10^6^ cells) were cotransfected with 2 μg of psPAX2-IN/HiBiT^63^, 2 μg of pWPI-Luc2^63^ and 1 μg of plasmids expressing SARS-CoV-2 S protein with Δ19CT using TransIT-LT1 according to the manufacturer’s protocol. After 2 days, cell culture supernatants were collected and filtered, and the pseudoviruses were stored at –80°C until use. The amount of pseudovirus prepared was quantified using the HiBiT assay. The HiBiT value measured by Nano Glo HiBiT lytic detection system (Promega, Cat# N3040) indicates the amount of p24 HIV-1 antigen. Then, the same amount of pseudoviruses normalized by the HiBiT value was inoculated into HOS-ACE2/TMPRSS2 cells (10,000 cells/well/50μl) in a 96 well plate. At 2 days postinfection, the infected cells were lysed with a Bright-Glo Luciferase Assay System (Promega, cat# E2620) and the luminescent signal was measured using a Centro XS3 LB960 plate reader (Berthhold Technologies) (**Fig. 4A, 4E**).

### BlaM-Vpr assay

HIV-1-based BlaM-Vpr assay^64,65^ was modified to perform SARS-CoV-2 S-based BlaM-Vpr assay. To produce SARS-CoV-2 S pseudovirus with BlaM-Vpr, HEK293T cells (2 × 10^6^ cells) were cotransfected with 1.6 μg of psPAX2-IN/HiBiT, 1.6 μg of pWPI-Luc2, 0.8 μg of plasmids expressing SARS-CoV-2 S protein with Δ19CT and 0.8 μg of BlaM-Vpr expression plasmid, pMM310 (HIV reagent program, Cat# ARP-11444) using TransIT-LT1 according to the manufacturer’s protocol. At 2 days posttransfection, cell culture supernatants were collected, filtered, and mixed with 40% polyethylene glycol #6000 (Nacalai Tesque Cat# 10200-25), followed by incubation at 4°C for 16 hours. After spinning down at 1,700 × g for 40 minutes at 4°C, the viral pellets were resuspended in 10%FBS/DMEM (high glucose) and the amount of pseudovirus prepared was quantified using the HiBiT assay. Note that the 24 well-collagen plate used in the following step should be kept at 4°C to inhibit starting fusion. The same amount of pseudoviruses normalized by HiBiT value was inoculated into HOS-ACE2/TMPRSS2 cells (2 × 10^5^ cells/well/1 ml) in a 24 well-collagen plate on ice and spinoculation was performed at 1,500 × g for 1 hour at 4°C to allow virus-cell attachment. After 1 hour, the cell culture supernatant including pseudoviruses was replaced with warmed fresh DMEM (high glucose), followed by incubation for 3 hours at 37°C. Then, each well was washed with phosphate-buffered saline (PBS) and β-lactamase loading solution (Invitrogen, Cat# K1030) [CCF4 substrate, solution B, solution C and anion transport inhibitor (Invitrogen, Cat# K1156) in Opti-MEM] was loaded. The plate was covered with aluminum foil to avoid light exposure and kept for 2 hours at room temperature. After 2 hours, cells were harvested, washed with PBS twice, and fixed with 2% paraformaldehyde phosphate (Nacalai Tesque, Cat# 09154-85). The fluorescence intensities at 520 nm (Pacific Blue; uncleaved CCF4) and 447 nm (AmCyan; cleaved CCF4) were detected by FACS Canto II (BD Biosciences) and analyzed by FlowJo software v10.7.1 (BD Biosciences). The ratio of cleaved CCF4 signal to the sum of uncleaved and cleaved CCF4 signals was calculated and shown as entry efficiency (**Fig. 5A, 5D**).

### SARS-CoV-2 infection

The day before infection, 1 × 10^4^ VeroE6/TMPRSS2 and Calu-3 cells were seeded into a 96-well plate. Then, VeroE6/TMPRSS2 (100 TCID_50_) and Calu-3 (5000 TCID_50_) cells were inoculated with SARS-CoV-2 and incubated at 37°C for 1 hour. After washing, 180 μl of fresh cell culture medium was added. 15 μl of cell culture supernatant was harvested at the indicated timepoints and used for RT-qPCR to quantify the viral RNA copy number (see “RT-qPCR” section) (**Fig. 6A, 6C**).

### RT-qPCR

RT-qPCR was performed as previously described^26,29,30,44,46^. Briefly, 5 μl of culture supernatant was mixed with 5 μl of 2 × RNA lysis buffer [2% Triton X-100 (Nacalai Tesque, Cat#12969-25), 50 mM KCl, 100mM Tris-HCl (pH 7.4), 40% glycerol, 0.8 U/μl recombinant RNase inhibitor (Takara, Cat# 2313A)] and incubated at room temperature for 10 minutes. 90 μl of RNase free water was added and then 2.5 μl of diluted sample was used for real-time RT-PCR according to the manufacturer’s protocol with One step TB green PrimeScript PLUS RT-PCR Kit (Takara, Cat# RR096A) and primers for *Nucleocapsid* (*N*) gene; Forward *N*, 5’-AGC CTC TTC TCG TTC CTC ATC-3’ and Reverse *N*, 5’-CCG CCA TTG CCA GCC ATT C-3’. The viral RNA copy number was standardized using a SARS-CoV-2 direct detection RT-qPCR kit (Takara, Cat# RC300A). Fluorescent signals from resulting PCR products were acquired using a Thermal Cycler Dice Real Time System III (Takara).

### Plaque assay

Plaque assay was performed as previously described^21,31,46,56^. One day before infection, 1 × 10^5^ VeroE6/TMPRSS2 cells were seeded into 24 well plate and infected with SARS-CoV-2 (50,000, 5000, 500, and 50 TCID_50_ respectively) at 37^◦^C for 1 hour. 3% FBS and 1.5% carboxymethyl cellulose (Wako, Cat# 039-1335) containing mounting solution was overlaid, followed by incubation at 37^◦^C. At 3 days postinfection, the cell culture medium was removed, and the cells were washed with PBS three times and fixed with 4% paraformaldehyde phosphate (Nacalai Tesque, Cat# 09154-85). The fixed cells were washed with tap water, dried, and stained with 0.1% methylene blue (Nacalai Tesque, Cat# 22412-14) in water for 30 minutes. The stained cells were washed with tap water and dried. The size of plaques was measured using Fiji software v2.2.0 (Image J) (**Fig. 3B**).

### Western blot analysis

As previously described, sample preparation for western blotting was performed with minor modifications^25,66–68^. The cell culture supernatants containing pseudoviruses produced from the HEK293T cells (see “Pseudovirus assay” section and “BlaM-Vpr assay” above) and the HEK293 cells transfected with the S expression plasmids (see “SARS-CoV-2 S-based fusion assay” section above) were used for western blotting. For viral lysate preparation, at 2 days posttransfection, cell culture supernatants were collected, filtered, and subjected to ultracentrifugation using 20% sucrose (22,000 × *g*, 4°C, 2 hours). Then, virions were dissolved in phosphate-buffered saline (PBS). To quantify HIV-1 p24 antigen in the pseudovirus, the amount of pseudoviruses in the cell culture supernatant was quantified by the HiBiT assay using a Nano Glo HiBiT lytic detection system. After normalization with HiBiT value, the samples were diluted with 2 × SDS sample buffer [100 mM Tris-HCl (pH6.8), 4% SDS, 12% β-mercaptoethanol, 20% glycerol, 0.05% bromophenol blue] and boiled for 5– 10 minutes at 100°C. For cell lysate preparation, the transfected cells were detached, washed twice with PBS, and lysed in lysis buffer [25mM HEPES (pH7.2), 20% glycerol, 125 mM NaCl, 1% Nonidet P40 substitute (Nacalai Tesque, Cat# 18558-54), protease inhibitor cocktail (Nacalai Tesque, Cat# 03969-21)]. Quantification of total protein in the cell lysates was done by protein assay dye (Bio-Rad, Cat# 5000006) according to manufacturer’s instruction. Then, cell lysates were diluted with 2 × SDS sample buffer and boiled for 5– 10 minutes. After cooling down, viral (pseudovirus) and cell lysates were mixed with diluted sample buffer (proteinsimple, Cat# 99351). Then, 5 × Fluorescent Master mix (proteinsimple, Cat# PS-ST01EZ-8) was added at a ratio of 4:1. Simple Western System was used for protein analysis. For protein detection, the following antibodies were used: rabbit anti-SARS-CoV-2 S (Novus Biologicals, Cat# NB100-56578, viral lysate; 1:40, cell lysate; 1:200). mouse anti-HIV-1 p24 monoclonal antibody (HIV Reagent Program, ARP-3537, 1:500), mouse anti-α tubulin monoclonal antibody (Sigma-Aldrich, Cat# T5168, 1:300), anti-rabbit secondary antibody (proteinsimple, Cat# 042-206), and anti-mouse secondary antibody (proteinsimple, Cat# 042-205). Bands were visualized and analyzed using Compass for Simple Western v6.1.0 (proteinsimple).

### Statistical analysis

Statistical significance was performed using two-sided unpaired or paired *t* test. GraphPad Prism software v8.4.3 (GraphPad Software) was used for these statistical tests unless otherwise mentioned. In the time-course experiments (**Figs. 1B, 6A, 6B**) a multiple regression analysis was performed to evaluate the statistical difference between experimental conditions thorough all timepoints. The initial time point was removed from the analysis. Familywise error rates (FWERs) were calculated by the Holm method. These analyses were performed in R v4.1.2 (https://www.r-project.org/).

## Acknowledgements

We would like to thank all members belonging to The Genotype to Phenotype Japan (G2P-Japan) Consortium. We thank Dr. Jin Gohda (The University of Tokyo, Japan) for providing reagents. We also thank Drs. Tomohiko Takasaki and Jun-Ichi Sakuragi (Kanagawa Prefectural Institute of Public Health) for providing SARS-CoV-2 Wuhan variant (strain SARS-CoV-2/Hu/DP/Kng/19-020, Genbank accession no. LC528232) and National Institute of Infectious Diseases for providing SARS-CoV-2 Alpha (strain QHN001, GISAID ID: EPI_ISL_804007), Beta (strain TY8-612, GISAID ID: EPI_ISL_1123289), Gamma (strain TY7-503, GISAID ID: EPI_ISL_877769), Delta (strain TY11-927, GISAID ID: EPI_ISL_2158617), Lambda (strain TY33-456, GISAID ID: EPI_ISL_4204973), Mu (strain TY26-717, GISAID ID: EPI_ISL_4470503), Omicron BA.1 (strain TY38-873, GISAID ID: EPI_ISL_7418017), and Omicron BA.2 (strain TY40-385, GISAID ID: EPI_ISL_9595859).

This study was supported in part by AMED SCARDA Japan Initiative for World-leading Vaccine Research and Development Centers “UTOPIA” (JP223fa627001, to Kei Sato), AMED SCARDA Program on R&D of new generation vaccine including new modality application (JP223fa727002, to Kei Sato); AMED Research Program on Emerging and Re-emerging Infectious Diseases (JP21fk0108574, to Hesham Nasser; JP22fk0108146, to Kei Sato; JP21fk0108494 to G2P-Japan Consortium, Terumasa Ikeda and Kei Sato; JP21fk0108425, to Kei Sato; 22fk0108506, Kei Sato; JP21fk0108432, to Kei Sato; JP22fk0108511, to G2P-Japan Consortium, Terumasa Ikeda and Kei Sato; JP22fk0108516, to Kei Sato; JP22fk0108506, to Kei Sato; JP22fk0108534, to Terumasa Ikeda and Kei Sato); AMED Research Program on HIV/AIDS (JP22fk0410055, to Terumasa Ikeda; and JP22fk0410039, to Kei Sato); JST CREST (JPMJCR20H4, to Kei Sato); JSPS KAKENHI Grant-in-Aid for Scientific Research C (21K07060, to Kenzo Tokunaga; 22K07103, to Terumasa Ikeda); JSPS KAKENHI Grant-in-Aid for Early-Career Scientists (22K16375, to Hesham Nasser); JSPS Core-to-Core Program (A. Advanced Research Networks) (JPJSCCA20190008, to Kei Sato); JSPS Leading Initiative for Excellent Young Researchers (LEADER) (to Terumasa Ikeda); The Cooperative Research Program (Joint Usage/Research Center program) of Institute for Life and Medical Sciences, Kyoto University (to Kei Sato); The Tokyo Biochemical Research Foundation (to Kei Sato); Takeda Science Foundation (to Terumasa Ikeda); Mochida Memorial Foundation for Medical and Pharmaceutical Research (to Terumasa Ikeda); The Naito Foundation (to Terumasa Ikeda); Waksman Foundation of Japan (to Terumasa Ikeda); an intramural grant from Kumamoto University COVID-19 Research Projects (AMABIE) (to Terumasa Ikeda); International Joint Research Project of the Institute of Medical Science, the University of Tokyo (to Terumasa Ikeda).

## The Genotype to Phenotype Japan (G2P-Japan) Consortium

Keita Matsuno^13^, Naganori Nao^13^, Hirofumi Sawa^13^, Shinya Tanaka^13^, Masumi Tsuda^13^, Lei Wang^13^, Yoshikata Oda^13^, Zannatul Ferdous^13^, Kenji Shishido^13^,Takasuke Fukuhara^13^, Tomokazu Tamura^13^, Rigel Suzuki^13^, Saori Suzuki^13^, Hayato Ito^13^, Jumpei Ito^6^, Yu Kaku^6^, Naoko Misawa^6^, Arnon Plianchaisuk^6^, Ziyi Guo^6^, Alfredo Jr. Hinay^6^, Keiya Uriu^6^, Yusuke Kosugi^6^, Shigeru Fujita^6^, Jarel Elgin Mendoza Tolentino^6^, Luo Chen^6^, Lin Pan^6^, Mai Suganami^6^, Mika Chiba^6^, Ryo Yoshimura^6^, Kyoko Yasuda^6^, Keiko Iida^6^, Naomi Ohsumi^6^, Adam Patrick Strange^6^, Hiroyuki Asakura^5^, Isao Yoshida^5^, So Nakagawa^14^, Akifumi Takaori-Kondo^15^, Kotaro Shirakawa^15^, Kayoko Nagata^15^, Ryosuke Nomura^15^, Yoshihito Horisawa^15^, Yusuke Tashiro^15^, Yugo Kawai^15^, Kazuo Takayama^15^, Rina Hashimoto^15^, Sayaka Deguchi^15^, Yukio Watanabe^15^, Ayaka Sakamoto^15^, Naoko Yasuhara^15^, Takao Hashiguchi^15^, Tateki Suzuki^15^, Kanako Kimura^15^, Jiei Sasaki^15^, Yukari Nakajima^15^, Hisano Yajima^15^, Takashi Irie^16^, Ryoko Kawabata^16^, Kaori Tabata^17^, Ryo Shimizu^1^, Yuka Mugita^1^, Takamasa Ueno^18^, Chihiro Motozono^18^, Mako Toyoda^18^, Akatsuki Saito^19^, Maya Shofa^19^, Yuki Shibatani^19^, Tomoko Nishiuchi^19^

^13^ Hokkaido University

^14^ Tokai University

^15^ Kyoto University

^16^ Hiroshima University

^17^ Kyushu University

^18^ Kumamoto University

^19^ Miyazaki University

## Contributions

MST Monira Begum, Kimiko Ichihara, Otowa Takahashi, Hesham Nasser, Michael Jonathan Kenzo Tokunaga, Kei Sato, and Terumasa Ikeda performed cell culture experiments.

Isao Yoshida, Mami Nagashima, Kenji Sadamasu, and Kazuhisa Yoshimura isolated clinical isolates and performed viral genome sequencing analysis.

Terumasa Ikeda designed the experiments and interpreted the results.

MST Monira Begum and Terumasa Ikeda wrote the original manuscript. All authors reviewed and proofread the manuscript.

The Genotype to Phenotype Japan (G2P-Japan) Consortium contributed to the project administration.

## Data availability

The raw data supporting the current study are available from the lead contact on request.

